# Broadly neutralizing antibodies against Omicron variants of SARS-CoV-2 derived from mRNA-lipid nanoparticle-immunized mice

**DOI:** 10.1101/2022.04.19.488843

**Authors:** Ruei-Min Lu, Kang-Hao Liang, Hsiao-Ling Chiang, Fu-Fei Hsu, Hsiu-Ting Lin, Wan-Yu Chen, Feng-Yi Ke, Monika Kumari, Yu-Chi Chou, Han-Chung Wu

**Author notes:** Corresponding author Dr. Han-Chung Wu Institute of Cellular and Organismic Biology, Academia Sinica No. 128, Academia Road, Section 2, Nankang, Taipei 11529, Taiwan.

## Abstract

The COVID-19 pandemic continues to threaten human health worldwide, as new variants of the severe acute respiratory syndrome coronavirus 2 (SARS-CoV-2) have emerged. Currently, the predominant circulating strains around the world are Omicron variants, which can evade many therapeutic antibodies. Thus, the development of new broadly neutralizing antibodies remains an urgent need. In this work, we address this need by using the mRNA-lipid nanoparticle immunization method to generate a set of Omicron-targeting monoclonal antibodies. Five of our novel K-RBD-mAbs show strong binding and neutralizing activities toward all SARS-CoV-2 variants of concern (Alpha, Beta, Gamma, Delta and Omicron). Notably, the epitopes of these five K-RBD-mAbs are overlapping and localized around K417 and F486 of the spike protein receptor binding domain (RBD). Chimeric derivatives of the five antibodies (K-RBD-chAbs) neutralize Omicron sublineages BA.1 and BA.2 with low IC_50_ values that range from 5.7 to 12.9 ng/mL. Additionally, we performed antibody humanization on a broadly neutralizing chimeric antibody to create K-RBD-hAb-62, which still retains excellent neutralizing activity against Omicron. Our results collectively suggest that these five therapeutic antibodies may effectively combat current and emerging SARS-CoV-2 variants, including Omicron BA.1 and BA.2. Therefore, the antibodies can potentially be used as universal neutralizing antibodies against SARS-CoV-2.

## Background

According to World Health Organization (WHO) reports through March 2022, more than 503 million people have been infected and 6.2 million have died as a result of the worldwide coronavirus disease 2019 (COVID-19) pandemic. The causal pathogen, severe acute respiratory syndrome coronavirus 2 (SARS-CoV-2), is a betacoronavirus with a linear single-stranded, positive-sense RNA genome (∼30 kb) encoding 29 proteins (Cao et al., 2021). SARS-CoV-2 infection is mediated by its spike protein, which is composed of an S2 domain and an S1 domain that contains an N-terminal domain (NTD), a C-terminal domain (CTD), and a receptor binding domain (RBD) (Walls et al., 2020). The RBD is an especially critical domain, as it initiates viral entry into host cells upon its binding to the host-cell receptor, angiotensin-converting enzyme 2 (ACE2). Due to its crucial role in infection, the RBD has been a common target for novel vaccines and neutralizing antibodies.

The high mutation rate associated with SARS-CoV-2 has contributed to its rapid global transmission, with many variants arising since the initial outbreak in late December 2019 (Harvey et al., 2021). Some of these variants are highly transmissible and can evade neutralization by vaccine-induced and therapeutic antibodies. Thus, the WHO has designated several “variants of concern” (VOCs), including Alpha (B.1.1.7), Beta (B.1.351), Gamma (P.1), Delta (B.617.2) and Omicron (B.1.1.529) (Hwang et al., 2022). Beta carries three mutations in the RBD (K417N, E484K and N501Y) and was found to effectively evade immune response and neutralizing antibodies (Wang et al., 2021). The Alpha variant became dominant globally in early 2021 due to its high transmissibility compared to previous lineages (Davies et al., 2021). In the third quarter of 2021, Alpha was gradually replaced by Delta, which exhibits even greater infectivity and pathogenicity, along with an ability to evade some neutralizing antibodies (Mlcochova et al., 2021; Sheikh et al., 2021).

On November 24, 2021, Omicron was first reported in South Africa, and two days later, WHO classified it as the fifth VOC. After just one month, Omicron had already outcompeted Delta to become the dominant circulating variant around the world; this rapid change is mainly due to the remarkably short caseload doubling time (2-3 days) (Grabowski et al., 2022; Jung et al., 2022). As of March 2022, the Omicron variant has diverged into four distinct sublineages: BA.1, BA.1.1, BA.2, and BA.3 (WHO, 2022). At the time of this writing, the most common circulating Omicron sublineages are BA.1 and BA.2. Compared to the original Wuhan-Hu-1 strain, Omicron BA.1 carries at least 33 mutations in the spike protein (29 amino acid substitutions, one insertion of three amino acids, and three small deletions), 15 of which are in the RBD. Ten of the 15 amino acid alterations in the RBD are located within the receptor-binding motif (RBM), which comprises the major site of contact with the human ACE2 receptor. The mutations in the RBM include: N440K, G446S, S477N, T478K, E484A, Q493K, G496S, Q498R, N501Y and Y505H. Together, these mutations increase the binding affinity of Omicron RBD to ACE2 by 2.4-fold and render the Omicron variants much more infectious than the Delta variant (Cameroni et al., 2022). Since the RBD is the primary target of neutralizing antibodies induced by COVID-19 vaccines, the accumulation of numerous mutations in this region has strongly reduced the affinity of vaccine-induced neutralizing antibodies, greatly weakening vaccine efficacy. Several studies have demonstrated that the current COVID-19 vaccines (i.e., mRNA-1273, BNT16b2, ChAdOx1 nCoV-19 and Ad26.COV2.S) have substantially reduced neutralizing potencies against Omicron, even in fully vaccinated individuals (Cameroni *et al*., 2022; Hoffmann et al., 2022; Liu et al., 2022; Planas et al., 2022).

Monoclonal antibodies (mAbs) have been widely used in basic research and clinical practice because of their high specificity and versatility (Lu et al., 2020). Such properties also make mAbs highly useful in the rapid development of antibody drugs and diagnostic kits to fight COVID-19 (Hwang *et al*., 2022). Over the past two year*s,* several neutralizing mAbs have shown *in vivo* efficacy for the prevention and treatment of SARS-CoV-2 (Corti et al., 2021; Taylor et al., 2021). For example, REGN-COV2 (combination of casirivimab and imdevimab) reduced the risk of COVID-19-related hospitalization and death by 70% in a cohort of non-hospitalized patients (Weinreich et al., 2021). REGN-COV2 was also able to prevent symptomatic COVID-19 and asymptomatic SARS-CoV-2 infection in previously uninfected household contacts of infected persons (O’Brien et al., 2021). Another neutralizing mAb treatment, AZD7442 (combination of tixagevimab and cilgavimab), caused a statistically significant reduction in the risk of developing symptomatic COVID-19, with protection from the virus continuing for at least 6 months (Loo et al., 2022). Altogether, the sustained efforts of researchers in academia and industry have so far yielded eight mAbs with emergency use authorizations (EUAs) granted by the U.S. FDA; these approved mAbs include: bamlanivimab, etesevimab, casirivimab, imdevimab, sotrovimab, cilgavimab, tixagevimab and bebtelovimab (Baum et al., 2020; Cathcart et al., 2022; Dong et al., 2021; Gottlieb et al., 2021; Shi et al., 2020; Westendorf et al., 2022). Unfortunately, pseudovirus and authentic virus experiments have shown that Omicron BA.1 is entirely resistant to bamlanivimab, etesevimab, casirivimab and imdevimab (IC_50_ >10 μg/mL), and it is partially resistant to sotrovimab, cilgavimab and tixagevimab (1 > IC_50_ > 0.1 μg/mL) (Cameroni *et al*., 2022; Cao et al., 2022; Dejnirattisai et al., 2022; Liu *et al*., 2022; Planas *et al*., 2022; Takashita et al., 2022a; VanBlargan et al., (IC_50_ < 0.01 μg/mL) (Westendorf *et al*., 2022). According to a recent study published in *Nature*, Omicron BA.1 can escape over 85% of 247 anti-RBD neutralizing antibodies in clinical use or under development (Cao *et al*., 2022).

In our previous work, we generated six chimeric neutralizing antibodies against the RBD of SARS-CoV-2 spike protein (Su et al., 2021). We showed that cocktails of these antibodies can neutralize D614G, Epsilon (B.1.429) and Kappa (B.1.617.1) strains, as well as four VOCs (Alpha, Beta, Gamma and Delta). The prophylactic and therapeutic activities of the individual neutralizing antibodies or antibody cocktails were then confirmed in D614G- and Delta-variant-infected mouse and hamster models (Liang et al., 2021). Although antibody therapy was effective in neutralizing the relevant VOCs of the time, we later found that Omicron BA.1 appears to be refractory to neutralization by our six mAbs. Since nearly all of the commercially available mAbs mentioned above are also ineffective against Omicron, there remains an unmet need for therapeutic antibodies with high neutralizing potency toward currently dominant variants. To quickly identify neutralizing antibodies for Omicron variants, we began by immunizing mice with messenger RNA (mRNA) encapsulated in lipid nanoparticles (LNPs), mimicking the now widely applied vaccine strategy. LNP-mediated delivery is a key aspect of the successful implementation of mRNA-based vaccines; BNT162b2 and mRNA-1273 each show >94% efficacy for preventing COVID-19 and have received EUAs in many countries (Baden et al., 2021; Hou et al., 2021; Polack et al., 2020). In our recent work, we established a platform that combines the mRNA-LNP immunization and hybridoma approaches, and we used it to rapidly generate neutralizing antibodies against the SARS-CoV-2 Delta variant (manuscript submitting). This time-saving method does not require manufacture of proteins as antigens, as the target protein is directly expressed in the animal and efficiently stimulates humoral immunity. In this study, we utilized bivalent mRNA-LNP encoding spike protein and the RBD of the Kappa variant to generate mAbs. The resulting mAbs were then screened for the ability to neutralize Omicron BA.1 and BA.2 pseudoviruses. Five of the most active anti-Omicron neutralizing antibodies were then engineered into chimeric formats and one was further humanized to enhance the potential for future use of these reagents in clinical settings.

## Materials and Methods

### Generation of mAbs using mRNA-LNP immunization method

An *in vitro* transcription system was used to generate mRNA encoding full-length spike and RBD of the SARS-CoV-2 Kappa variant (QTZ19256.1); the mRNA also contained a 5’ UTR, IgG kappa leader sequence, 3’ UTR and poly(A) tail. Individual lipids were dissolved in ethanol and mixed. DLin-MC3-DMA (MC3) (MedChemExpress), 1, 2-distearoyl-sn-glycero-3-phosphocholine (DSPC) (Avanti Polar Lipids), cholesterol (Sigma-Aldrich) and PEG-2000 (MedChemExpress) were combined at molar ratios of 50:10:38.5:1.5. The lipid mixture was then combined with a 50 mM sodium acetate buffer pH 4.5, containing mRNA at a ratio of 3:1 (aqueous: ethanol), prior to mixing in NanoAssemblr® IGNITE NxGen Cartridges (Precision NanoSystems Inc.). After formulation, LNPs were dialyzed against PBS (pH 7.4) and concentrated using Amicon Ultra-Centrifugal Filter with a 10-kDa cutoff (Merck).

Four- to six-week-old female BALB/c mice were immunized with 5 μg of the Kappa spike and RBD mRNA-LNP by intramuscular (I.M.) injection. After four inoculations with the same concentration of mRNA-LNP, the splenocytes from immunized mice were harvested and fused with mouse myeloma NS-1 cells. The fused cells were cultured in DMEM supplemented with 15% FBS, HAT medium, and hybridoma cloning factors in 96-well tissue culture plates (Lu et al., 2019). Two weeks after fusion, the culture supernatants were screened by ELISA. Selected clones were subcloned via limiting dilutions. Hybridoma clones were isotyped using a commercially available ELISA isotyping it (Southern Biotech, Birmingham, AL, USA). All animal experiments were approved by the Academia Sinica Institutional Animal Care and Use Committee (IACUC protocol No. 20051468).

### Recombinant protein-based ELISA

Recombinant RBD and spike-His tag proteins for different SARS-CoV-2 variants were purchased from ACROBiosystems. ELISA plates were coated with 0.5 μg/mL recombinant protein in 0.1 M NaHCO_3_ (pH 8.6) buffer at 4°C overnight, followed by blocking with PBS containing 1% bovine serum albumin (BSA) at room temperature for 2 h. After blocking, the wells were washed twice with PBS. Anti-RBD and control antibodies were added to the plates and incubated for 1 h at room temperature. The plates were washed with PBS containing 0.1% Tween-20 (PBST_0.1_) three times and then incubated for 1 h with Peroxidase-conjugated secondary antibody (Jackson ImmunoResearch). After three washes with PBST_0.1_, signal was produced using 3,3’5,5’-Tetramethylbenzidine (TMB) substrate (TMBW-1000-01, SURMODICS). The reaction was stopped with 3 N HCl, and absorbance was measured at 450 nm with an ELISA plate reader.

### Pseudovirus neutralization assay

Pseudovirus expressing SARS-CoV-2 spike protein was provided by the National RNAi Core Facility (Academia Sinica, Taiwan). The pseudovirus neutralization assays were performed using HEK293T cells that expressed human ACE2 (HEK293T/hACE2) seeded on 96-well white plates (SPL Life Science) at a density of 1 × 10^4^ cells per well. Serial dilutions of K-RBD-mAbs were pre-incubated with 1000 TU SARS-CoV-2 pseudovirus in 1% FBS DMEM for 1 h at 37°C. The mixtures were then added to pre-seeded HEK293T/hACE2 cells for 24 h at 37°C. The pseudovirus-containing culture medium was removed and replaced with 10% FBS DMEM for an additional 48-h incubation. Next, ONE-Glo luciferase reagent (Promega) was added to each well for a 3-min incubation at 37°C. The luminescence was measured with a microplate spectrophotometer (ID3, Molecular Devices). The half maximal inhibitory concentration (IC_50_) was calculated by nonlinear regression using Prism software version 8.1.0 (GraphPad Software Inc.).

### Identification of V_H_ and V_K_ sequence of mAbs

Total RNA was extracted from hybridoma cells using TRIzol reagent (Thermo Scientific), and cDNA was generated with an oligo(dT)_20_ primer and Superscript III reverse transcriptase (Thermo Scientific). The V_H_ and V_L_ gene fragments were amplified from the cDNA by PCR using primer sets directed against the mouse immunoglobulin (Ig) variable region (Zhou et al., 1994). The PCR products were cloned using the pGEM-T Easy Vector System (Promega) and analyzed by DNA sequencing. From the sequences, the framework regions (FRs) and complementarity determining regions (CDRs) were defined by searching with the NCBI IgBLAST program (https://www.ncbi.nlm.nih.gov/igblast/).

### Cellular ELISA

The binding of K-RBD-mAbs to the RBD mutants was examined by cellular ELISA. Human HEK293T cells were transiently transfected in 6-well plates with wild-type or mutant RBD plasmids. The next day, transfected cells were seeded on 96-well plates. The cells were fixed with 4% paraformaldehyde in PBS for 15 min at room temperature at 48 h post-transfection and then incubated in 0.1% triton X-100 at room temperature for 10 min. After blocking the cells with 5% milk, antibodies against RBD were added to the wells (100 ng/mL) for 1 h at room temperature. Then, the cells were washed and horseradish peroxidase-conjugated anti-human antibody (1:2000) was added for 1 h at room temperature. Signal was generated with TMB and was measured on an ELISA plate reader.

### Construction and expression of chimeric antibodies (chAbs)

The V_H_ and V_K_ gene segments of mAbs were amplified by PCR and introduced via appropriate restriction enzyme sites. The V_H_ genes were cloned in-frame into a modified expression vector with a signal peptide and human IgG1 constant region. The V_K_ genes were cloned into a modified expression vector with a signal peptide and human kappa chain constant region. The heavy-chain- and light-chain-encoding plasmids were co-transfected into Expi-293F cells (Thermo Scientific) to produce antibodies. Protein G column chromatography (GE healthcare) was performed to purify chimeric IgGs. After dialysis of eluents with PBS, the antibody concentrations were measured using the bicinchoninicacid (BCA) protein assay (Thermo Scientific).

### Antibody humanization

A CDR grafting approach was used to generate the humanized antibody from K-RBD-mAb-62. First, the amino acid sequence of the V_H_ and V_L_ domains of the mAb were aligned with the human Ig variable domain germline database using the NCBI IgBLAST tool. The IGHV1-2*06 and IGKV1-33*01 sequences were selected as the most similar human Ig sequences. The V_H_ and V_L_ of the mAb were then respectively grafted onto the V_H_ and V_L_ frameworks of the selected human Ig gene. Both genes were synthesized and amplified by PCR using Kapa-Hifi DNA polymerase (Roche). The resulting V_H_ was cloned into a modified pcDNA5-FRT-Gamma1 expression vector with human IgG_1_ constant region. The V_L_ was cloned into a modified p-Kappa-HuGS expression vector. A homologous 3D structure of the Fab was then built based on a previously described computer modeling method (Waterhouse et al., 2018). After analyzing the structure with PyMOL software, we identified the key amino acid residues at which mutations may impact the original conformation of the CDRs. These residues were back-mutated to the corresponding mouse residues.

## Results

### Generation and Identification of mAbs against SARS-CoV-2 Spike Protein

To generate novel mAbs that bind to SARS-CoV-2 spike protein, we immunized BALB/c mice with LNP-encapsulated mRNAs encoding Kappa spike and RBD. Sixty-one mAbs against Kappa RBD (K-RBD-mAbs) were generated using the mouse hybridoma technique, according to ELISA-based mAb reactivity measurements (Table 1). Most K-RBD-mAbs also recognized the Wuhan and Delta RBDs. Importantly, 27 mAbs in the hybridoma culture supernatants could cross-react with Omicron BA.1 RBD and spike protein. We therefore purified the IgGs of these clones and assessed their binding to Omicron BA.1 RBD and spike (Fig.1A and B). K-RBD-mAb-60, −62, −75 and −111 had the highest binding activities for both target proteins. Next, we further used SARS-CoV-2 spike-pseudotyped lentivirus to screen the neutralizing potentials of the 27 Omicron BA.1-cross-reactive K-RBD-mAbs (Fig. 1C). Among the strongest Omicron spike-binding antibodies, K-RBD-mAb-60, −62, and −75, showed high neutralizing activity, whereas K-RBD-mAb-111 did not. In addition, K-RBD-mAb-42 and −19 had only moderate binding activities but exhibited strong neutralizing activities. Of note, the antibody isotypes of the five neutralizing mAbs were IgG2a/kappa (Table 1).

**Figure 1.**
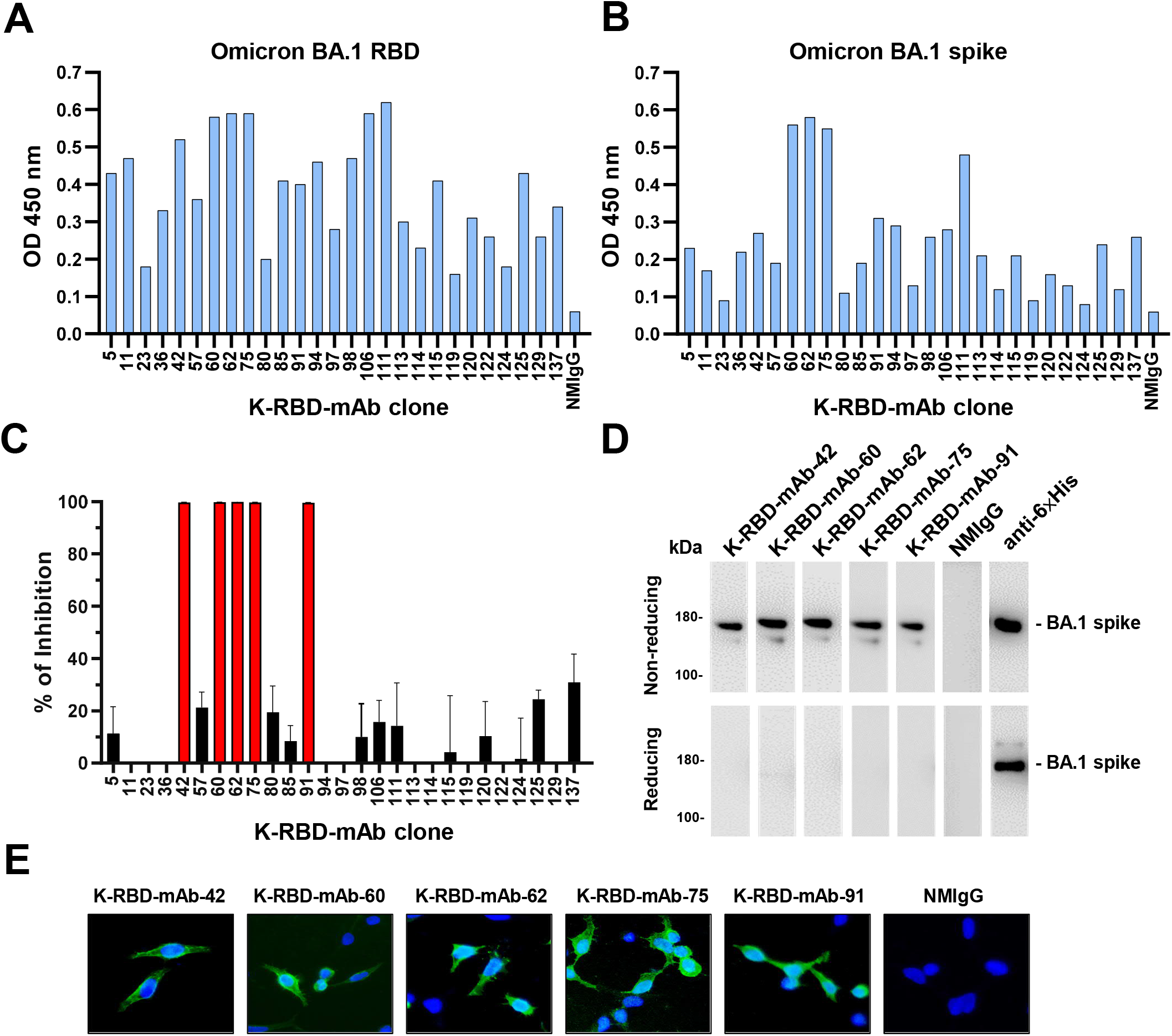
Identification of mAbs targeting the RBD of the Omicron BA.1 variant. Comparative ELISA was performed to assess binding activity of 27 K-RBD-mAbs toward the BA.1 RBD (A) and trimeric BA.1 spike protein (B). ELISA plates were coated with recombinant BA.1 RBD-His and trimeric spike protein (2 μg/mL) and incubated with mAbs that were diluted to 500 ng/mL. Normal mouse IgG (NMIgG) was used as a negative control. (C) Neutralization assay of 27 K-RBD-mAbs (1 μg/mL) for BA.1. Data represent one of two independent experiments. (D) Five K-RBD-mAbs were used as primary antibodies for Western blotting of recombinant BA.1 spike protein-His. Anti-6×His mAb was used as a positive control. (E) BA.1 spike-expressing 293T cells were probed with 1 μg/mL K-RBD-mAbs and then stained with FITC goat anti-mouse IgG. NMIgG was used as a negative control.

**Table 1.**
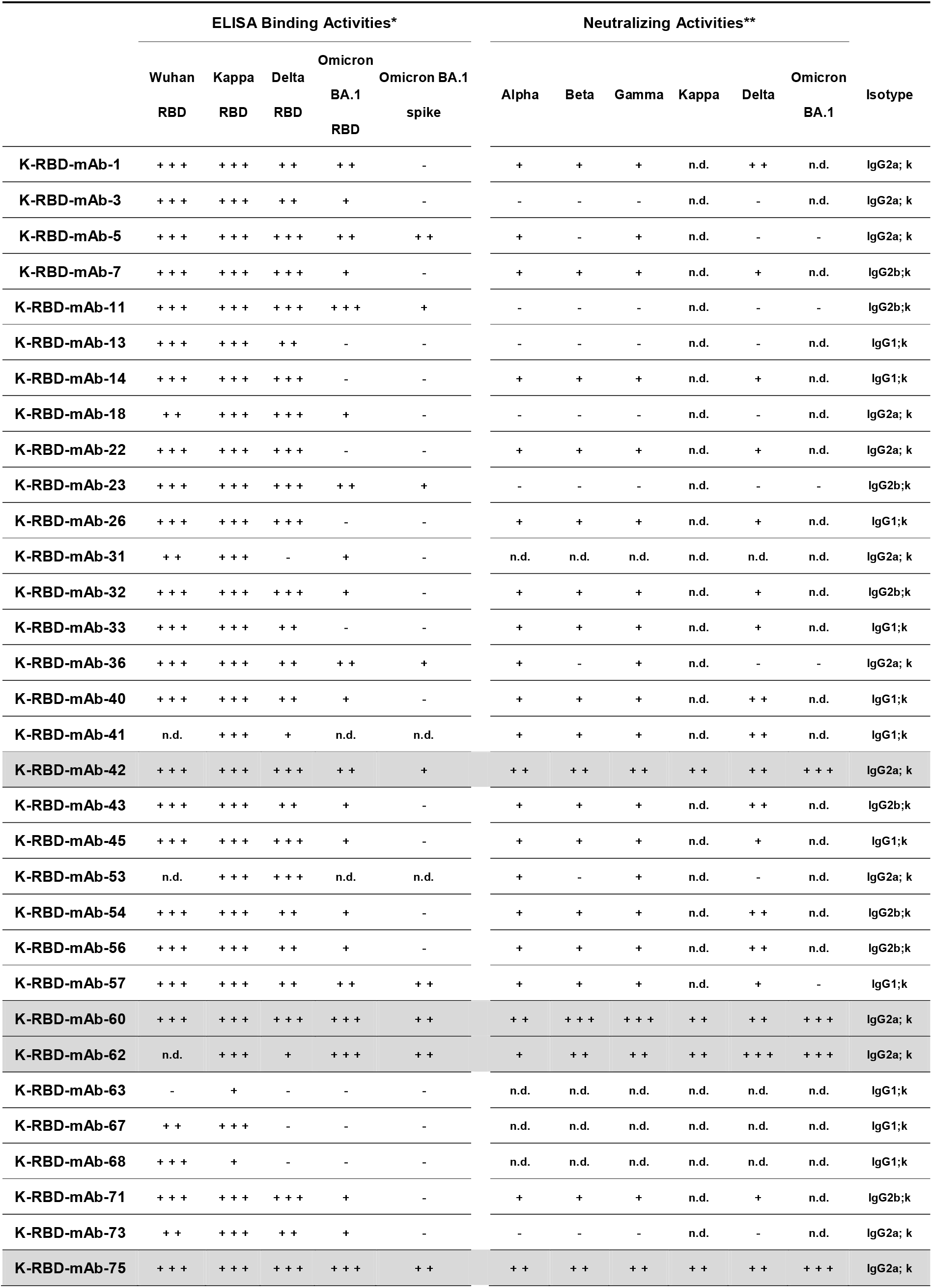
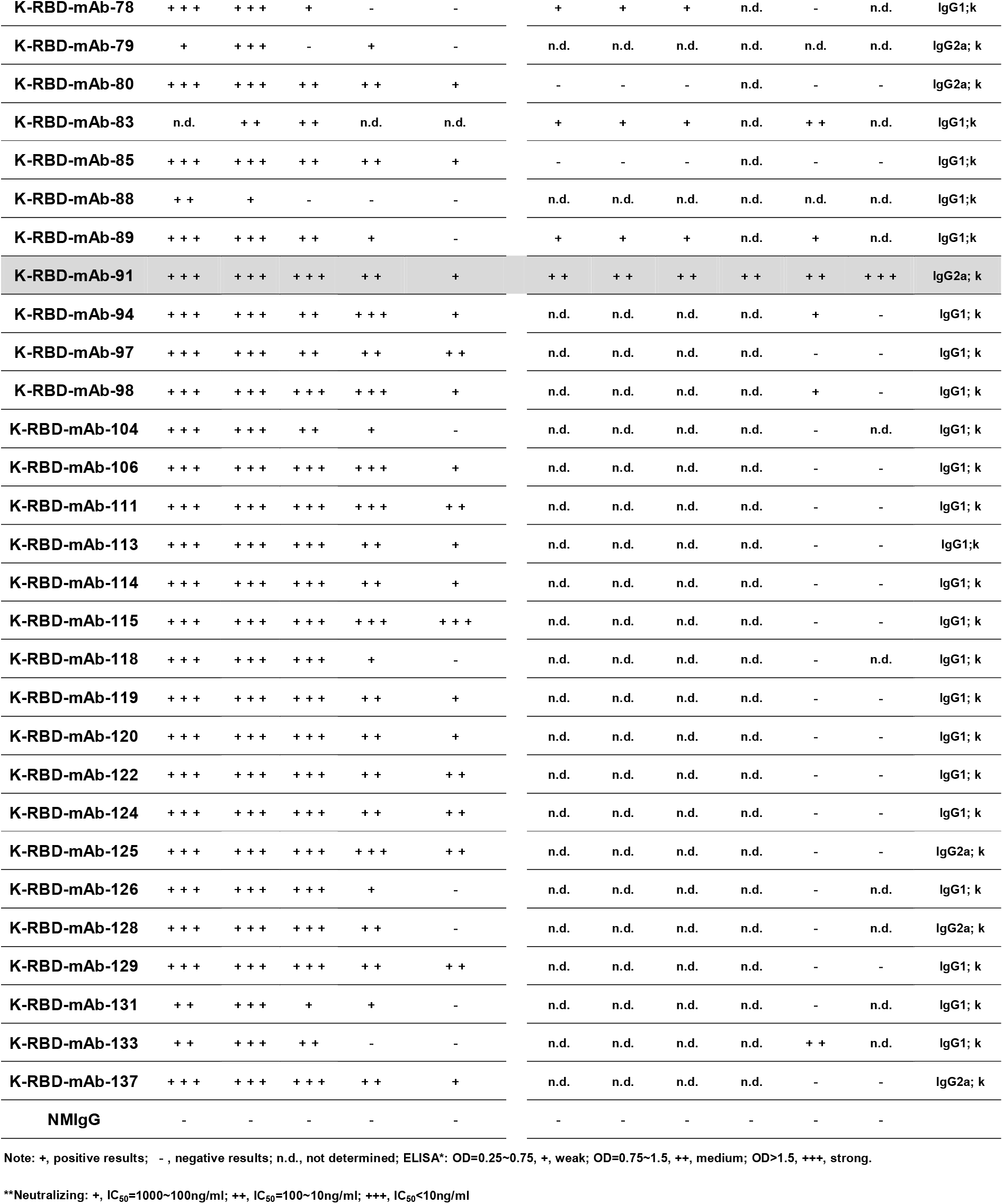
Characterization of the mAbs activities against RBD of SARS-CoV-2.

All five Omicron BA.1 neutralizing antibodies (K-RBD-mAb-19, −−2, −60, −62 and −75) could recognize recombinant Omicron BA.1 spike protein at its expected molecular weight of 180 kDa in a Western blot analysis under non-reducing conditions (Fig. 1D). The five K-RBD-mAbs could also be used to stain Omicron BA.1 spike protein-expressing 293T cells but not mock-transfected cells, indicating that these mAbs can be applied in immunofluorescence experiments (Fig. 1E).

### Neutralizing abilities of K-anti-RBD mAbs

We examined the neutralizing abilities of five K-RBD-mAbs toward SARS-CoV-2 Kappa and five VOCs, including Alpha, Beta, Gamma, Delta and Omicron BA.1 (Fig. 2A). Pseudovirus neutralization assays revealed that all five K-RBD-mAbs were capable of neutralizing all six variants, and each mAb exhibited high neutralizing activity toward Omicron BA.1 (IC_50_ values ranged between 3.6 to 7.1 ng/mL; Fig. 2B). Among the five mAbs, K-RBD-mAb-62 showed the lowest IC_50_ value (7.0 ng/mL) toward the Delta variant, and K-RBD-mAb-60 exhibited the best neutralizing abilities toward the Alpha, Beta, Gamma and Omicron BA.1 variant. Overall, the K-RBD-mAbs had broad neutralizing activities against SARS-CoV-2 variants, including Omicron BA.1.

**Figure 2.**
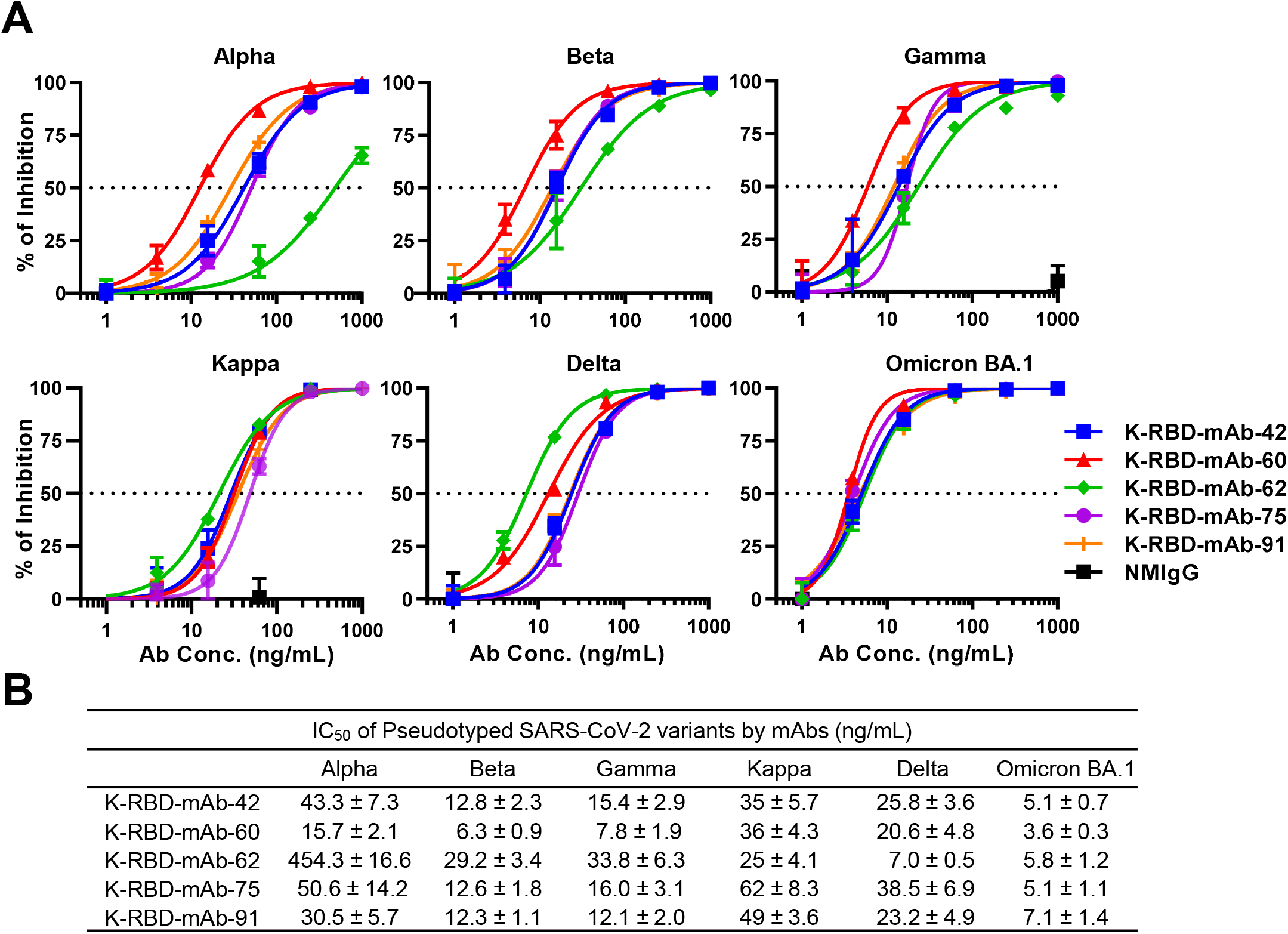
Neutralizing capacities of K-RBD-mAbs toward SARS-CoV-2 variant pseudoviruses. (A) Dose-response of K-RBD-mAbs in a neutralization assay of SARS-CoV-2 variant pseudovirus. Data are the mean ± SE of three independent experiments. (B) For each antibody, the presented IC_50_ value was calculated with GraphPad Prism software.

### The binding epitopes of K-anti-RBD mAbs

To investigate whether these mAbs share overlapping epitopes, we carried out ELISA-based competition assays (Fig. 3A). P44 is a non-neutralizing anti-BA.1 RBD mAb, so it was used as a non-competitor control. We found no positive ELISA signals in competition assays between the five K-RBD-mAbs, suggesting that each mAb fully competes with the others for binding to the RBD. These results suggested that the epitopes for all five mAb mostly overlap. Since the RBD residues, K417, Y453, Q474, F486, Q498, T500 and N501, are the amino acid residues that directly contact ACE2 (Yan et al., 2020), we sought to clarify whether any of these residues are targeted by the five K-RBD-mAbs. To do so, we mutated each residue to alanine and transiently expressed each mutant in 293T cells. We then performed alanine scanning by cellular ELISA to test the impacts of each mutation on binding of the K-RBD-mAbs (Fig. 3B). The results showed that singleton mutations at Y453 and F486 dramatically decreased the binding of all five mAbs (by more than 80%). The mutation at K417 moderately reduced the binding of the five mAbs. In contrast to the other mAbs, K-RBD-mAb-62 was more sensitive to the Q474 mutation. The combinatorial mutations of K417A/Y453A and Q474A/F486A both substantially disrupted the binding of K-RBD-mAbs (Fig. 3C). The combined Q498A/T500A/N501A mutations also had a similar effect on K-RBD-mAbs. Taken together, these data suggest that the RBD epitopes recognized by all five K-RBD-mAbs are very similar.

**Figure 3.**
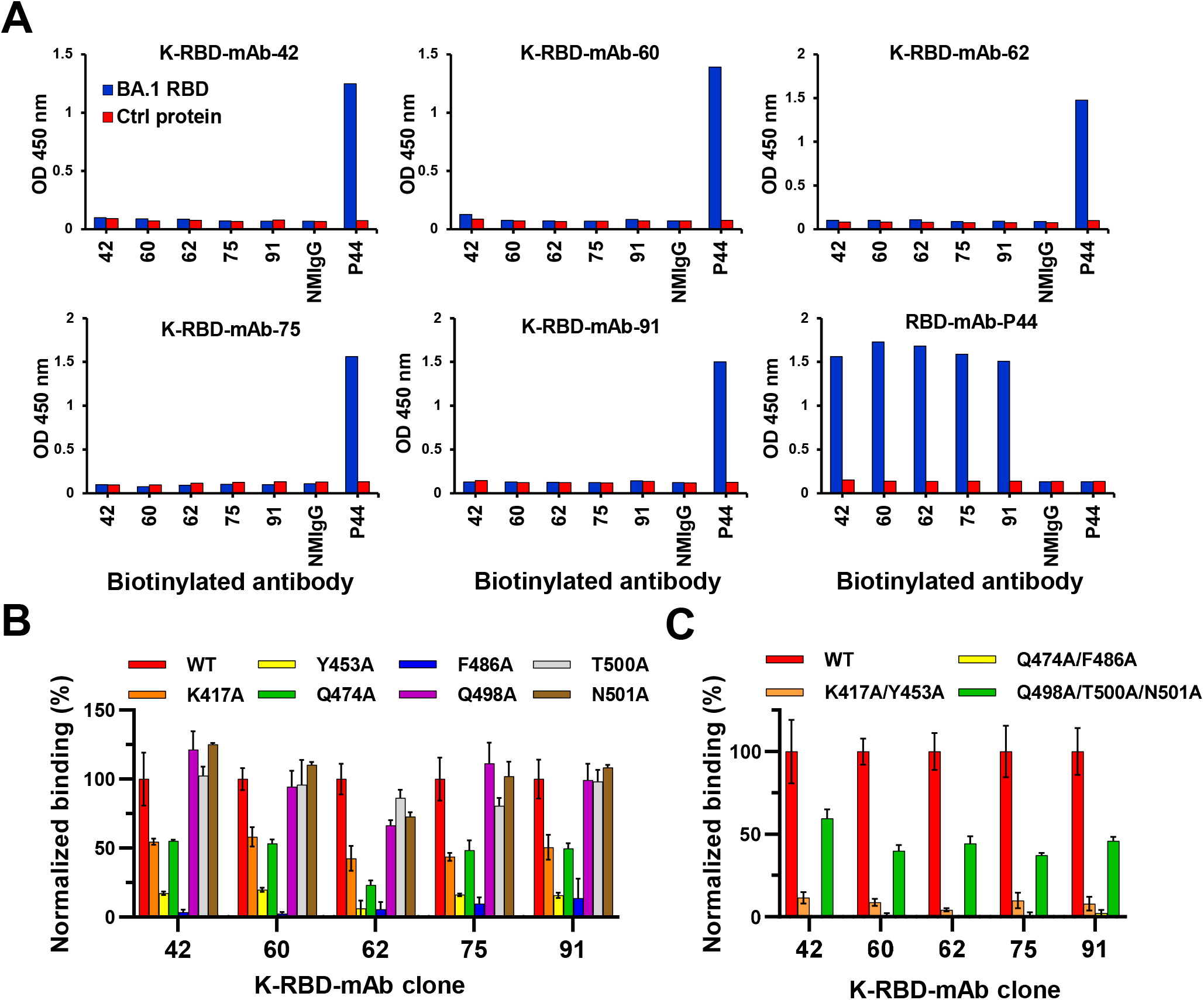
Epitope mapping of neutralizing K-RBD-mAbs. In the ELISA-based epitope competition assay, BA.1 RBD protein was captured by K-RBD-mAb that was immobilized on a 96-well plate. Then, biotinylated antibodies were added to compete for binding of BA.1 RBD. HRP-conjugated streptavidin was added to detect the biotinylated antibodies. Non-competitively binding antibodies produced absorbance signals at OD 450 nm. EpEX-His served as a negative control protein (Ctrl). P44, a non-neutralizing mAb against BA.1 RBD, was used as a positive control. (B-C) Epitope mapping of K-RBD-mAbs according to the mutagenesis assay. 293T cells were made to transiently express wild-type (WT) or mutant RBD proteins with single or multiple alanine mutations. Binding of each mAb to the RBD mutants was examined by cellular ELISA. The results were normalized and are presented as a percent control.

### Generation of K-RBD chimeric antibodies (chAbs)

To improve the potential of these mAbs for clinical use, we identified the V_H_ and V_L_ genes of the five K-RBD mAbs and respectively grafted them onto human IgG1 and kappa backbones to generate K-RBD-chAbs. Notably, the ELISA results showed all five K-RBD-chAbs not only bound to the RBD and spike protein of Omicron BA.1 but also to that of BA.2 (Fig. 4A). K-RBD-chAb-60 showed the highest binding activities when tested against the BA.1 and BA.2 spike proteins. We next assessed the neutralization profiles of the five K-RBD-chAbs against the variants of SARS-CoV-2 (Fig. 4B). The five chAbs exerted substantial neutralizing activities against BA.1 and BA.2 pseudoviruses, with IC_50_ values ranging from 5.7 to 12.9 ng/mL (Fig. 4C).

**Figure 4.**
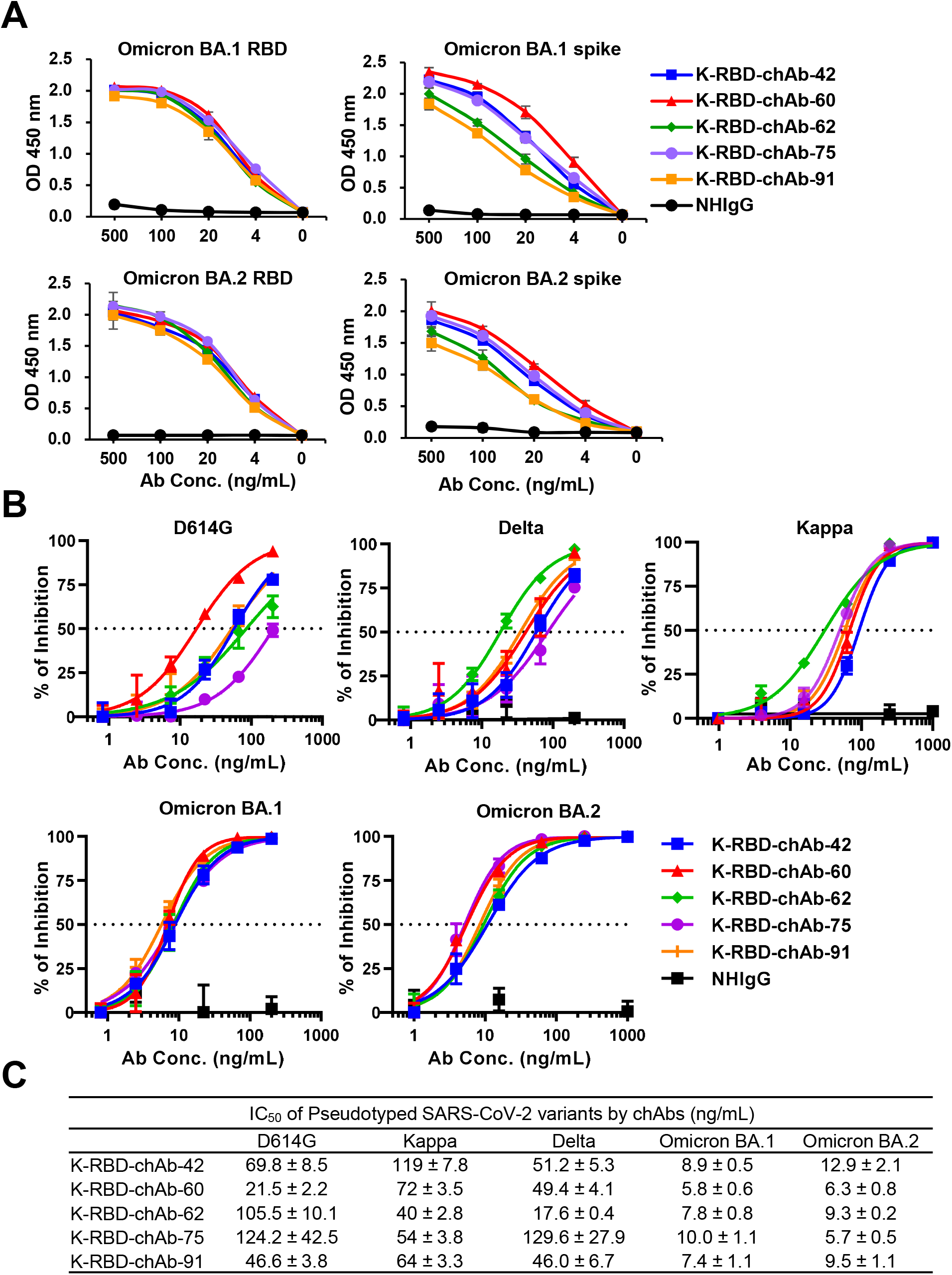
Generation of Omicron-neutralizing K-RBD chAbs. (A) ELISA was used to measure binding activity of five K-RBD chAbs to RBD and spike proteins of Omicron BA.1 and BA.2. Serial dilutions were made to evaluate each chAb at 500 to 0 ng/mL. (B) Individual neutralization curves for K-RBD-chAbs were generated based on pseudovirus neutralization assays. Error bars denote the mean ± SE for three technical replicates. (C) For each antibody, the IC_50_ value was calculated with GraphPad Prism software from three independent experiments, and the values are presented in the table.

### Synthesis of Humanized Therapeutic Antibody

The IC_50_ of K-RBD-chA-62 for Omicron is less than 10 ng/mL, and its IC_50_ values toward Kappa and Delta are the lowest of the five chAbs (Fig. 4C). Hence, we chose to further develop K-RBD-chA-62 by synthesizing a humanized version using CDR grafting techniques. The resulting humanized antibody is named K-RBD-hAb-62, and its amino acid sequence shares ∼93% identity with human IgG1/Kappa. Since the binding activity of K-RBD-hAb-62 against the RBDs of BA.1 and BA.2 appeared to be identical to its chimeric format (Fig. 5A), it is likely that the humanization process did not greatly affect its paratope. Finally, we demonstrated that K-RBD-hAb-62 neutralized BA.1 and BA.2 with respective IC_50_ values of 7.0 and 9.5 ng/mL (Figure 5B), indicating that its Omicron-neutralizing activities were retained.

**Figure 5.**
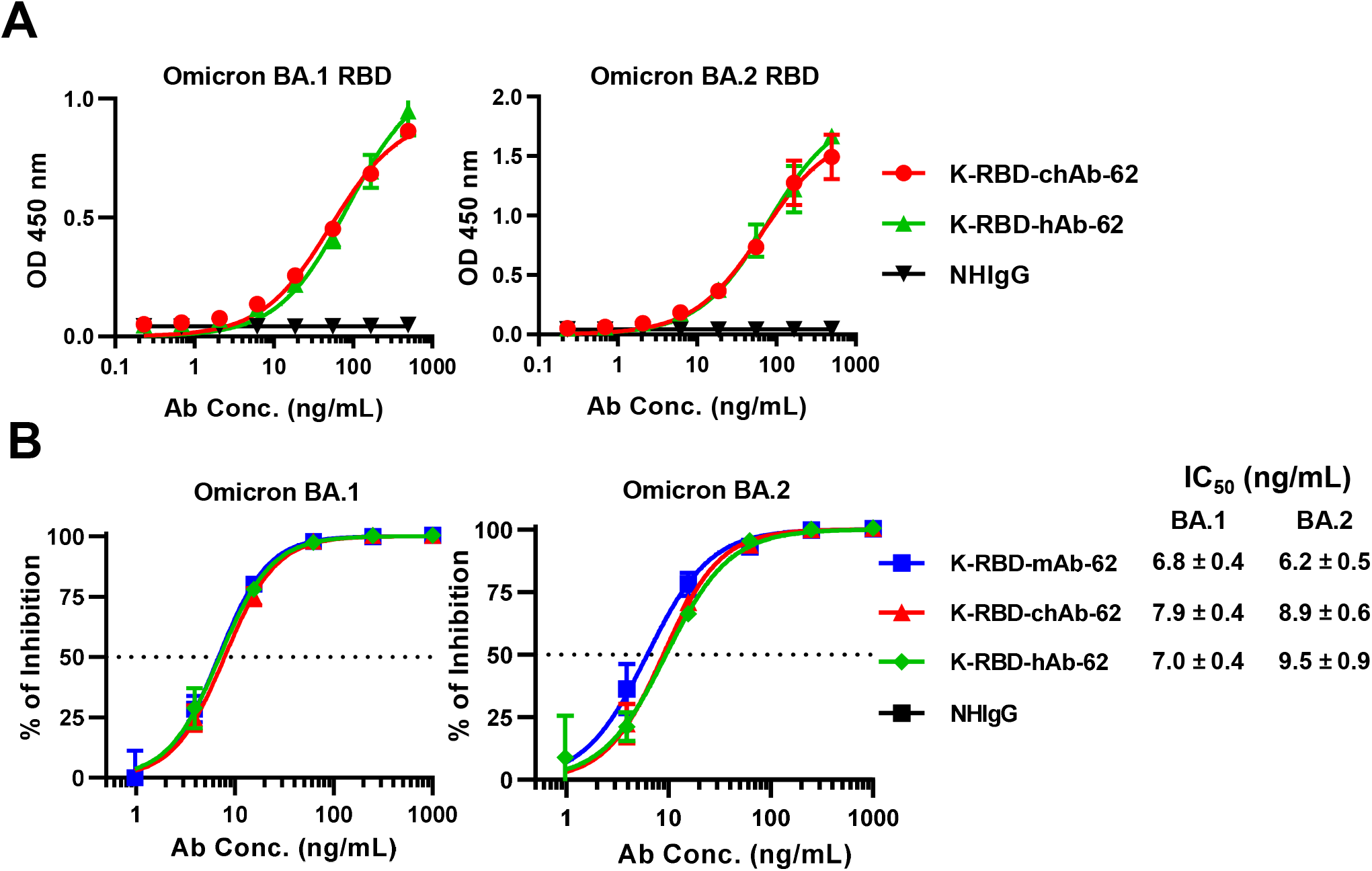
Generation of Omicron-neutralizing K-RBD humanized antibody. (A) Omicron binding activities of K-RBD-hAb-62 and K-RBD-chAb-62 were determined by ELISA and compared. (B) Neutralization of Omicron BA.1 and BA.2 pseudovirus by three versions of K-RBD-mAb-62. The IC_50_ value was calculated with Prism software. Each assay was performed in three independent experiments. All data points are shown, along with the mean ± SE.

## Discussion

Four major strategies are often used to generate therapeutic mAbs: mouse hybridoma, phage-display Ab library, hAb transgenic mice, and single B cell RT-PCR (Lu *et al*., 2020). The most common approach for rapid isolation of neutralizing antibodies against SARS-CoV-2 has been single cell RT-PCR on memory B cells sorted from convalescent or acute-phase COVID-19 patient samples (Liu et al., 2020; Rogers et al., 2020). In addition, the classical mouse hybridoma approach plus further Ab engineering techniques have also contributed to the discovery of neutralizing antibodies during the COVID-19 pandemic (Su *et al*., 2021). In this study, we used a modified mouse hybridoma technique including mRNA-LNP immunization method to generate mAbs that can broadly neutralize SARS-CoV-2 variant pseudotypes. To the best of our knowledge, only one patent application from Novatis (Dominy et al., 2018) has so far described the use of this efficient method to generate mAbs.

The high number of mutations in the Omicron spike protein have allowed the variant to evade neutralizing antibodies elicited by vaccines and prior natural infections (Liu *et al*., 2022; Planas *et al*., 2022). This immune evasion may be one of the major reasons that Omicron variants have spread so much more quickly than the Wuhan-Hu-1 virus and the Delta variant. Furthermore, nearly all of the therapeutic Abs granted EUAs have been severely compromised by the emergence of Omicron BA.1 (Dejnirattisai *et al*., 2022; VanBlargan *et al*., 2022). For example, bamlanivimab binds the right shoulder of the RBD and is sensitive to changes at E484. Beta, Gamma and Omicron variants all contain mutations at this site and cannot be neutralized by bamlanivimab. The epitopes of etesevimab overlap with the binding site of ACE2, and its binding is reduced by RBD mutations at K417, D420, F456, A475 and L455. Tixagevimab and casirivimab were found to be very sensitive to changes at F486, N487 and G476 in the RBD, and the multiple mutations S477N/T478K/E484A in Omicron BA.1 significantly decrease the neutralizing activities of these two antibodies. Imdevimab and cilgavimab target the loop-forming residues 440-449 in the RBD; thus, the N440K, G446S, S477N mutations in Omicron BA.1 greatly reduce the neutralization activities of both antibodies. G339 is a part of the sotrovimab epitope, and the G339D mutation in Omicron BA.1 reduces the IC_50_ value of sotrovimab to 200 ng/mL (Cameroni *et al*., 2022). Here we showed K417, Y453, Q474 and F486 might be common residues in the epitopes of our five neutralizing K-RBD-mAbs (Fig. 3B). Interestingly, the K417N, S477N and E484A mutations in Omicron BA.1 and BA.2 are near the putative epitopes of our K-RBD-mAbs, but the mutations did not affect the binding and neutralization activities of these antibodies (Fig. 4). In future work, we plan to better define the binding interface of K-RBD-mAbs and Omicron RBD by utilizing cryo-electron microscopy (cryo-EM) analyses.

The BA.2 and BA.1 variants share twelve common mutations in the RBD, while BA.2 has four additional mutations: S371F, T376A, D405N and R408S (Yu et al., 2022). In recent months, the number of BA.2 cases has rapidly increased according to the GISAID database (https://covidcg.org), making it the dominant strain in many countries and suggesting that BA.2 has a selective advantage over BA.1. Using the live-virus focus reduction neutralization test (FRNT), Takashita et al. found that etesevimab and bamlanivimab lost neutralizing activity against BA.2 and BA.1 (Takashita et al., 2022b). Interestingly, imdevimab (REGN10987), retained neutralizing activity against BA.2 (68.7 ng/mL of FRNT _50_), though it was previously shown to have lost neutralizing activity against BA.1. Notably, this report also showed that the FRNT_50_ value for sotrovimab against BA.2 was 1359 ng/mL, which represents a 49.7-fold increase over the value for the wild-type strain (Takashita *et al*., 2022b). According to new data included in the health care provider fact sheet, the neutralization of BA.2 pseudovirus and authentic virus by sotrovimab was significantly weaker than that of wild-type virus, with a 16-fold reduction in IC_50_ value (GlaxoSmithKline, 2022). The U.S. FDA announced that the authorized dosage of sotrovimab is unlikely to be effective against the BA.2 variant, and the agency updated the EUA to limit sotrovimab use in states where BA.2 is dominant such as New Jersey and New York (FDA, 2022). Fortunately, the IC_50_ value of bebtelovimab for BA.2 neutralization was not changed, and this treatment is still expected to be effective against the BA.2 variant (FDA, 2022). Because of the high mutation rate in the SARS-CoV-2 RNA genome, extensive and continual development of therapeutic antibodies is needed, and combinations of existing neutralizing antibodies should be evaluated for the ability to control new variants of SARS-CoV-2.

The advantages of using mRNA vaccines over traditional protein-based technologies are beginning to be defined. The manufacture of recombinant proteins requires sophisticated technologies and is expensive, resulting in long lead times. Alternatively, mRNA has cost advantages and can be synthesized by a generic process once the nucleic acid sequence is known (Hou *et al*., 2021; Van Hoecke and Roose, 2019). mRNA technology also has some advantages related to its mode of *in vivo* protein expression; the use of exogenous mRNA expression can overcome significant barriers to the application of shortLlived proteins, but it may have no impact on treatment duration for long lived proteins (Karikó et al., 2012; Thess et al., 2015). In addition to its remarkable potential as a vaccine substrate, the broad applicability of mRNA technology has been demonstrated in the field of passive immunotherapy (antibody therapeutics) (Schlake et al., 2019). A previous study showed that single injections of LNPLencapsulated mRNAs encoding heavy chain and light chain can rapidly stimulate expression of anti-rabies and botulinum mAbs *in vitro* and *in vivo*, thereby conferring prophylactic and therapeutic protection in mice (Thran et al., 2017). In another study, humanized anti-HER2 Ab (trastuzumab) mRNA was formulated into LNPs for efficient *in vivo* delivery, and the anticancer activity of this treatment was demonstrated (Rybakova et al., 2019). Furthermore, CD5-targeting LNPs were used to deliver mRNA encoding anti-fibroblast activation protein (FAP) chimeric antigen receptor and generate anti-FAP CAR-T cells *in vivo* (Rurik et al., 2022). The resulting CAR-T cells successfully reduced fibrosis and restored cardiac function after injury. Here, we used our recently established mRNA-LNP-mediated immunization method to generate mAbs against infectious diseases, which may further expand the spectrum of mRNA applications.

In conclusion, we used our mRNA-LNP immunization platform to generate anti-RBD mAbs that bind to and neutralize the five current SARS-CoV-2 VOCs. The engineered K-RBD-chAbs and K-RBD-hAb-62 neutralized Omicron sublineages BA.1 and BA.2 with low IC_50_ values. Thus, these neutralizing antibodies may be promising tools for controlling current SARS-CoV-2 variants, including Omicron.

## List of abbreviations

COVID-19: Coronavirus disease 2019
SARS-CoV-2: severe acute respiratory syndrome coronavirus 2
RBD: receptor-binding domain
RBM: receptor binding motif
VOC: variant of concern
chAb: chimeric antibody
hAb: humanized antibody;
IC_50_: half maximal inhibitory concentration
EUA: emergency use authorization
FDA: Food and Drug Administration

## Declarations

Not applicable.

## Consent for publication

Not applicable.

## Availability of supporting data

All materials and supporting data will be made available upon reasonable request to the corresponding author.

## Authors’ contributions

RML designed experiments, wrote the manuscript and performed antibody humanization. WYC and HTL immunized the mice and generated mouse mAbs. KHL and HLC performed *in vitro* neutralization by pseudoviruses. HTL and WYC performed neutralizing Ab expression. FFH and MK synthesized mRNA-LNP. YCC generated pseudoviruses. HCW conceived the experiments, obtained funding, wrote and revised the paper, and provided overall direction for the study.

## Acknowledgments

Academia Sinica, for their assistance with Ab production and analysis. We also thank the National RNAi Core Facility at BioTReC for providing the SARS-CoV-2 variant pseudoviruses.

## References

Baden, L.R., El Sahly, H.M., Essink, B., Kotloff, K., Frey, S., Novak, R., Diemert, D., Spector, S.A., Rouphael, N., and Creech, C.B. (2021). Efficacy and safety of the mRNA-1273 SARS-CoV-2 vaccine. New England journal of medicine 384, 14. 10.1056/NEJMoa2035389.

Baum, A., Fulton, B.O., Wloga, E., Copin, R., Pascal, K.E., Russo, V., Giordano, S., Lanza, K., Negron, N., and Ni, M. (2020). Antibody cocktail to SARS-CoV-2 spike protein prevents rapid mutational escape seen with individual antibodies. Science 369, 1014–1018.

Cameroni, E., Bowen, J.E., Rosen, L.E., Saliba, C., Zepeda, S.K., Culap, K., Pinto, D., VanBlargan, L.A., De Marco, A., di Iulio, J., et al. (2022). Broadly neutralizing antibodies overcome SARS-CoV-2 Omicron antigenic shift. Nature 602, 664–670. 10.1038/s41586-021-04386-2.

Cao, C., Cai, Z., Xiao, X., Rao, J., Chen, J., Hu, N., Yang, M., Xing, X., Wang, Y., Li, M., et al. (2021). The architecture of the SARS-CoV-2 RNA genome inside virion. Nature Communications 12, 3917. 10.1038/s41467-021-22785-x.

Cao, Y., Wang, J., Jian, F., Xiao, T., Song, W., Yisimayi, A., Huang, W., Li, Q., Wang, P., An, R., et al. (2022). Omicron escapes the majority of existing SARS-CoV-2 neutralizing antibodies. Nature 602, 657–663. 10.1038/s41586-021-04385-3.

Cathcart, A.L., Havenar-Daughton, C., Lempp, F.A., Ma, D., Schmid, M.A., Agostini, M.L., Guarino, B., Di iulio, J., Rosen, L.E., Tucker, H., et al. (2022). The dual function monoclonal antibodies VIR-7831 and VIR-7832 demonstrate potent in vitro and in vivo activity against SARS-CoV-2. bioRxiv, 2021.2003.2009.434607. 10.1101/2021.03.09.434607.

Corti, D., Purcell, L.A., Snell, G., and Veesler, D. (2021). Tackling COVID-19 with neutralizing monoclonal antibodies. Cell 184, 3086–3108.

Davies, N.G., Abbott, S., Barnard, R.C., Jarvis, C.I., Kucharski, A.J., Munday, J.D., Pearson, C.A.B., Russell, T.W., Tully, D.C., Washburne, A.D., et al. (2021). Estimated transmissibility and impact of SARS-CoV-2 lineage B.1.1.7 in England. Science 372, eabg3055. doi:10.1126/science.abg3055.

Dejnirattisai, W., Huo, J., Zhou, D., Zahradník, J., Supasa, P., Liu, C., Duyvesteyn, H.M.E., Ginn, H.M., Mentzer, A.J., Tuekprakhon, A., et al. (2022). SARS-CoV-2 Omicron-B.1.1.529 leads to widespread escape from neutralizing antibody responses. Cell 185, 467-484.e415. https://doi.org/10.1016/j.cell.2021.12.046.

Dominy, J., Dunn, R., Glaser, S., Keating, M., Klattenhoff, C., and Splawski, I. (2018). Mrna-mediated immunization methods. PCT patent application WO2018/029586, 04.08.2017.

Dong, J., Zost, S.J., Greaney, A.J., Starr, T.N., Dingens, A.S., Chen, E.C., Chen, R.E., Case, J.B., Sutton, R.E., Gilchuk, P., et al. (2021). Genetic and structural basis for SARS-CoV-2 variant neutralization by a two-antibody cocktail. Nature microbiology 6, 1233–1244. 10.1038/s41564-021-00972-2.

FDA (2022). FDA updates Sotrovimab emergency use authorization. https://www.fda.gov/drugs/drug-safety-and-availability/fda-updates-sotrovimab-emergency-use-authorization.

GlaxoSmithKline (2022). Sotrovimab Fact Sheet for Healthcare Providers. https://www.sotrovimab.com/.

Gottlieb, R.L., Nirula, A., Chen, P., Boscia, J., Heller, B., Morris, J., Huhn, G., Cardona, J., Mocherla, B., and Stosor, V. (2021). Effect of bamlanivimab as monotherapy or in combination with etesevimab on viral load in patients with mild to moderate COVID-19: a randomized clinical trial. JAMA 325, 632–644.

Grabowski, F., Kochańczyk, M., and Lipniacki, T. (2022). The Spread of SARS-CoV-2 Variant Omicron with a Doubling Time of 2.0–3.3 Days Can Be Explained by Immune Evasion. Viruses 14, 294.

Harvey, W.T., Carabelli, A.M., Jackson, B., Gupta, R.K., Thomson, E.C., Harrison, E.M., Ludden, C., Reeve, R., Rambaut, A., Peacock, S.J., et al. (2021). SARS-CoV-2 variants, spike mutations and immune escape. Nature Reviews Microbiology 19, 409–424. 10.1038/s41579-021-00573-0.

Hoffmann, M., Krüger, N., Schulz, S., Cossmann, A., Rocha, C., Kempf, A., Nehlmeier, I., Graichen, L., Moldenhauer, A.-S., Winkler, M.S., et al. (2022). The Omicron variant is highly resistant against antibody-mediated neutralization: Implications for control of the COVID-19 pandemic. Cell 185, 447–456.e411. https://doi.org/10.1016/j.cell.2021.12.032.

Hou, X., Zaks, T., Langer, R., and Dong, Y. (2021). Lipid nanoparticles for mRNA delivery. Nature Reviews Materials 6, 1078–1094. 10.1038/s41578-021-00358-0.

Hwang, Y.C., Lu, R.M., Su, S.C., Chiang, P.Y., Ko, S.H., Ke, F.Y., Liang, K.H., Hsieh, T.Y., and Wu, H.C. (2022). Monoclonal antibodies for COVID-19 therapy and SARS-CoV-2 detection. Journal of Biomedical Science 29, 1. 10.1186/s12929-021-00784-w.

Jung, C., Kmiec, D., Koepke, L., Zech, F., Jacob, T., Sparrer, K.M.J., and Kirchhoff, F. (2022). Omicron: what makes the latest SARS-CoV-2 variant of concern so concerning? Journal of virology, jvi0207721. 10.1128/jvi.02077-21.

Karikó, K., Muramatsu, H., Keller, J.M., and Weissman, D. (2012). Increased Erythropoiesis in Mice Injected With Submicrogram Quantities of Pseudouridine-containing mRNA Encoding Erythropoietin. Molecular Therapy 20, 948–953. https://doi.org/10.1038/mt.2012.7.

Liang, K.H., Chiang, P.Y., Ko, S.H., Chou, Y.C., Lu, R.M., Lin, H.T., Chen, W.Y., Lin, Y.L., Tao, M.H., Jan, J.T., and Wu, H.C. (2021). Antibody cocktail effective against variants of SARS-CoV-2. Journal of Biomedical Science 28, 80. 10.1186/s12929-021-00777-9.

Liu, L., Iketani, S., Guo, Y., Chan, J.F.W., Wang, M., Liu, L., Luo, Y., Chu, H., Huang, Y., Nair, M.S., et al. (2022). Striking antibody evasion manifested by the Omicron variant of SARS-CoV-2. Nature 602, 676–681. 10.1038/s41586-021-04388-0.

Liu, L., Wang, P., Nair, M.S., Yu, J., Rapp, M., Wang, Q., Luo, Y., Chan, J.F., Sahi, V., Figueroa, A., et al. (2020). Potent neutralizing antibodies against multiple epitopes on SARS-CoV-2 spike. Nature 584, 450–456. 10.1038/s41586-020-2571-7.

Loo, Y.M., McTamney, P.M., Arends, R.H., Abram, M.E., Aksyuk, A.A., Diallo, S., Flores, D.J., Kelly, E.J., Ren, K., and Roque, R. (2022). The SARS-CoV-2 monoclonal antibody combination, AZD7442, is protective in non-human primates and has an extended half-life in humans. Science Translational Medicine, eabl8124.

Lu, R.M., Chiu, C.Y., Liu, I.J., Chang, Y.L., Liu, Y.J., and Wu, H.C. (2019). Novel human Ab against vascular endothelial growth factor receptor 2 shows therapeutic potential for leukemia and prostate cancer. Cancer Sci 110, 3773–3787. 10.1111/cas.14208.

Lu, R.M., Hwang, Y.C., Liu, I.J., Lee, C.C., Tsai, H.Z., Li, H.J., and Wu, H.C. (2020). Development of therapeutic antibodies for the treatment of diseases. Journal of Biomedical Science 27, 1. 10.1186/s12929-019-0592-z.

Mlcochova, P., Kemp, S.A., Dhar, M.S., Papa, G., Meng, B., Ferreira, I.A., Datir, R., Collier, D.A., Albecka, A., and Singh, S. (2021). SARS-CoV-2 B. 1.617. 2 Delta variant replication and immune evasion. Nature 599, 114-119.

O’Brien, M.P., Forleo-Neto, E., Musser, B.J., Isa, F., Chan, K.C., Sarkar, N., Bar, K.J., Barnabas, R.V., Barouch, D.H., Cohen, M.S., et al. (2021). Subcutaneous REGEN-COV Antibody Combination to Prevent Covid-19. The New England journal of medicine 385, 1184–1195. 10.1056/NEJMoa2109682.

Planas, D., Saunders, N., Maes, P., Guivel-Benhassine, F., Planchais, C., Buchrieser, J., Bolland, W.-H., Porrot, F., Staropoli, I., Lemoine, F., et al. (2022). Considerable escape of SARS-CoV-2 Omicron to antibody neutralization. Nature 602, 671–675. 10.1038/s41586-021-04389-z.

Polack, F.P., Thomas, S.J., Kitchin, N., Absalon, J., Gurtman, A., Lockhart, S., Perez, J.L., Pérez Marc, G., Moreira, E.D., Zerbini, C., et al. (2020). Safety and Efficacy of the BNT162b2 mRNA Covid-19 Vaccine. New England Journal of Medicine 383, 2603–2615. 10.1056/NEJMoa2034577.

Rogers, T.F., Zhao, F., Huang, D., Beutler, N., Burns, A., He, W.T., Limbo, O., Smith, C., Song, G., Woehl, J., et al. (2020). Isolation of potent SARS-CoV-2 neutralizing antibodies and protection from disease in a small animal model. Science 369, 956–963. 10.1126/science.abc7520.

Rurik, J.G., Tombácz, I., Yadegari, A., Fernández, P.O.M., Shewale, S.V., Li, L., Kimura, T., Soliman, O.Y., Papp, T.E., Tam, Y.K., et al. (2022). CAR T cells produced in vivo to treat cardiac injury. Science 375, 91–96. doi:10.1126/science.abm0594.

Rybakova, Y., Kowalski, P.S., Huang, Y., Gonzalez, J.T., Heartlein, M.W., DeRosa, F., Delcassian, D., and Anderson, D.G. (2019). mRNA Delivery for Therapeutic Anti-HER2 Antibody Expression In Vivo. Molecular Therapy 27, 1415–1423. 10.1016/j.ymthe.2019.05.012.

Schlake, T., Thess, A., Thran, M., and Jordan, I. (2019). mRNA as novel technology for passive immunotherapy. Cellular and Molecular Life Sciences 76, 301–328. 10.1007/s00018-018-2935-4.

Sheikh, A., McMenamin, J., Taylor, B., and Robertson, C. (2021). SARS-CoV-2 Delta VOC in Scotland: demographics, risk of hospital admission, and vaccine effectiveness. The Lancet 397, 2461–2462.

Shi, R., Shan, C., Duan, X., Chen, Z., Liu, P., Song, J., Song, T., Bi, X., Han, C., Wu, L., et al. (2020). A human neutralizing antibody targets the receptor-binding site of SARS-CoV-2. Nature 584, 120–124. 10.1038/s41586-020-2381-y.

Su, S.C., Yang, T.J., Yu, P.Y., Liang, K.H., Chen, W.Y., Yang, C.W., Lin, H.T., Wang, M.J., Lu, R.M., Tso, H.C., et al. (2021). Structure-guided antibody cocktail for prevention and treatment of COVID-19. PLOS Pathogens 17, e1009704. 10.1371/journal.ppat.1009704.

Takashita, E., Kinoshita, N., Yamayoshi, S., Sakai-Tagawa, Y., Fujisaki, S., Ito, M., Iwatsuki-Horimoto, K., Chiba, S., Halfmann, P., Nagai, H., et al. (2022a). Efficacy of Antibodies and Antiviral Drugs against Covid-19 Omicron Variant. New England Journal of Medicine 386, 995–998. 10.1056/NEJMc2119407.

Takashita, E., Kinoshita, N., Yamayoshi, S., Sakai-Tagawa, Y., Fujisaki, S., Ito, M., Iwatsuki-Horimoto, K., Halfmann, P., Watanabe, S., Maeda, K., et al. (2022b). Efficacy of Antiviral Agents against the SARS-CoV-2 Omicron Subvariant BA.2. New England Journal of Medicine, https://www.nejm.org/doi/full/10.1056/NEJMc2201933.

Taylor, P.C., Adams, A.C., Hufford, M.M., de la Torre, I., Winthrop, K., and Gottlieb, R.L. (2021). Neutralizing monoclonal antibodies for treatment of COVID-19. Nature reviews. Immunology 21, 382–393. 10.1038/s41577-021-00542-x.

Thess, A., Grund, S., Mui, B.L., Hope, M.J., Baumhof, P., Fotin-Mleczek, M., and Schlake, T. (2015). Sequence-engineered mRNA Without Chemical Nucleoside Modifications Enables an Effective Protein Therapy in Large Animals. Molecular Therapy 23, 1456–1464. https://doi.org/10.1038/mt.2015.103.

Thran, M., Mukherjee, J., Pönisch, M., Fiedler, K., Thess, A., Mui, B.L., Hope, M.J., Tam, Y.K., Horscroft, N., Heidenreich, R., et al. (2017). mRNA mediates passive vaccination against infectious agents, toxins, and tumors. EMBO molecular medicine 9, 1434–1447. 10.15252/emmm.201707678.

Van Hoecke, L., and Roose, K. (2019). How mRNA therapeutics are entering the monoclonal antibody field. Journal of Translational Medicine 17, 54. 10.1186/s12967-019-1804-8.

VanBlargan, L.A., Errico, J.M., Halfmann, P.J., Zost, S.J., Crowe, J.E., Purcell, L.A., Kawaoka, Y., Corti, D., Fremont, D.H., and Diamond, M.S. (2022). An infectious SARS-CoV-2 B.1.1.529 Omicron virus escapes neutralization by therapeutic monoclonal antibodies. Nature Medicine 28, 6. 10.1038/s41591-021-01678-y.

Walls, A.C., Park, Y.-J., Tortorici, M.A., Wall, A., McGuire, A.T., and Veesler, D. (2020). Structure, function, and antigenicity of the SARS-CoV-2 spike glycoprotein. Cell 181, 281–292. e286.

Wang, P., Nair, M.S., Liu, L., Iketani, S., Luo, Y., Guo, Y., Wang, M., Yu, J., Zhang, B., and Kwong, P.D. (2021). Antibody resistance of SARS-CoV-2 variants B. 1.351 and B. 1.1. 7. Nature 593, 130-135.

Waterhouse, A., Bertoni, M., Bienert, S., Studer, G., Tauriello, G., Gumienny, R., Heer, F.T., de Beer, T.A.P., Rempfer, C., Bordoli, L., et al. (2018). SWISS-MODEL: homology modelling of protein structures and complexes. Nucleic acids research 46, 8. 10.1093/nar/gky427.

Weinreich, D.M., Sivapalasingam, S., Norton, T., Ali, S., Gao, H., Bhore, R., Xiao, J., Hooper, A.T., Hamilton, J.D., Musser, B.J., et al. (2021). REGEN-COV Antibody Combination and Outcomes in Outpatients with Covid-19. New England Journal of Medicine 385, e81. 10.1056/NEJMoa2108163.

Westendorf, K., Wang, L., Žentelis, S., Foster, D., Vaillancourt, P., Wiggin, M., Lovett, E., van der Lee, R., Hendle, J., Pustilnik, A., et al. (2022). LY-CoV1404 (bebtelovimab) potently neutralizes SARS-CoV-2 variants. bioRxiv, https://doi.org/10.1101/2021.1104.1130.442182.

WHO (2022). Weekly epidemiological update on COVID-19. https://www.who.int/emergencies/diseases/novel-coronavirus-2019/situation-reports.

Yan, R., Zhang, Y., Li, Y., Xia, L., Guo, Y., and Zhou, Q. (2020). Structural basis for the recognition of SARS-CoV-2 by full-length human ACE2. Science 367, 1444–1448. 10.1126/science.abb2762.

Yu, J., Collier, A.-r.Y., Rowe, M., Mardas, F., Ventura, J.D., Wan, H., Miller, J., Powers, O., Chung, B., Siamatu, M., et al. (2022). Neutralization of the SARS-CoV-2 Omicron BA.1 and BA.2 Variants. New England Journal of Medicine, https://www.nejm.org/doi/full/10.1056/NEJMc2201849.

Zhou, H., Fisher, R.J., and Papas, T.S. (1994). Optimization of primer sequences for mouse scFv repertoire display library construction. Nucleic Acids Research 22, 888–889. 10.1093/nar/22.5.888.

